# Ancestral Quadruplication of Protein Kinase R in Tarsiers Hints Towards Novel Primate-Virus Innovations

**DOI:** 10.64898/2026.10.23.761390

**Authors:** Padraic G Heneghan, Tristan R Grieve, Meru J Sadhu

## Abstract

Tarsiers, an elusive family of primates found in Indonesia and the Philippines, sit at an essential point in primate evolution, with their divergence between that of Strepsirrhini (lemurs) and Simiiformes (monkeys and apes). Through homology searches, we have discovered that tarsiers carry the first known case in primates of a gene expansion of protein kinase R (PKR), a kinase involved in the response to viral infection. PKR was quadruplicated in the ancestor of tarsiers and has been maintained since then in all sequenced tarsier species. The extra copies lack the double-stranded RNA (dsRNA) binding domains necessary for detecting viral infection yet largely maintain the residues necessary for the kinase function that carries out PKR’s antiviral function. We show that three of the four PKR copies indeed maintain kinase activity, one of which appears to be resistant to the viral PKR antagonist K3 from vaccinia virus. Our analyses of site and branch-site positive selection reveal that the tarsier PKRs are under positive selection. We propose that these extra copies could serve as a diversification strategy to bypass viral inhibitors that bind PKR’s kinase domain, or they could alternatively serve as decoys that shield the canonical PKR from viral inhibition. This is the first instance of mammal PKR duplicates lacking dsRNA-binding domains and may highlight an innovation in an ancient, long-standing molecular arms race between viruses and a close relative of humans.

## Introduction

Protein kinase R (PKR), also called elongation initiation factor 2 alpha kinase 2 (eIF2AK2), is one of the four kinases that triggers the integrated stress response (ISR). The ISR is essential in halting cell translation under stressful conditions, such as hypoxia, nutrient deprivation, and viral infection. PKR is specifically involved in the detection of viral infection for this pathway (reviewed in (Pakos-Zebrucka, et al. 2016)).

PKR is a multi-domain protein composed of two double-stranded RNA binding domains (dsRBDs) at its N-terminus, linked to a serine/threonine kinase at its C-terminus (Zhang, et al. 2001). During infection, viruses can produce double-stranded RNA (dsRNA) during viral transcription. Binding to viral dsRNA by the dsRBDs facilitates PKR dimerization, leading to autophosphorylation and activation of its kinase domain; the active kinase then proceeds to phosphorylate a serine residue (S51) on eIF2ɑ, ceasing translation in the cell to block viral protein production (Zhang, et al. 2001; Dar, et al. 2005). This makes PKR a key component of the anti-viral innate immune system.

PKR is the fastest evolving of the four kinases in the ISR (HRI, PERK, and GCN2), and it is hypothesized this is specifically due to its role in interacting with viruses (Elde, et al. 2009; Rothenburg, et al. 2009). Viruses have a variety of strategies for evading PKR. In some cases, viruses encode proteins that directly bind and inhibit PKR; for instance, K3 is an effector encoded by several poxviruses as a mimic of eIF2ɑ, competing for PKR binding to block eIF2ɑ phosphorylation and, if successful, permitting viral infection (Kawagishi-Kobayashi, et al. 1997; Elde, et al. 2009). Given the shared interface of eIF2ɑ and K3, PKR and K3 are a popular pair in the research of molecular arms races, whereby PKR must make substitutions to evade K3 while maintaining binding to eIF2ɑ (Chambers, et al. 2024; Chambers, et al. 2026). PKR has many residues under positive selection, some of which have been attributed to evading viral mimics like K3 (Elde, et al. 2009; Rothenburg, et al. 2009).

Genes with roles in innate immunity sometimes expand into gene families, which could be a way to defend against diverse pathogens or counter pathogen evasion (Daugherty, et al. 2016; Tenthorey, et al. 2022). Expansions of PKR have been noted in frogs and fish, including the PKZ gene, which is derived from PKR but has the dsRNA binding domains swapped for Z-conformation nucleic acid-binding domains (Rothenburg, et al. 2008). PKR has mostly been found as a single copy in mammalian genomes (Wu, et al. 2024). However, very recent telomere-to-telomere assemblies of bat genomes have revealed several *Myotis* species have haplotype variation in copy number of *PKR*, and that some of these duplicate PKR proteins are resistant to viral mimics (Jacquet, et al. 2022; Vazquez, et al. 2026). In these bats, there are haplotypes of both expanded and single-copy *PKR* genes, implying the selective pressure of viral conflict remains to maintain the additional copies (Vazquez, et al. 2026).

Here, we report the first evidence of a *PKR* gene expansion in a family of primates, the Tarsiidae, common-name tarsiers. This is the second example of *PKR* expansions in a family of mammals. Tarsiers were only sequenced in the last decade, despite being at a key point in primate evolution, diverging between the emergence of the Strepsirrhini (lemurs, galagos, lorises, and pottos) and Catarrhini and Platyrrhini (Old World and New World monkeys) (Schmitz, et al. 2016). We demonstrate that the PKR expansion is maintained across the four sequenced species of the family, meaning it is quite ancient. We suspect that the maintenance of this expansion may be due to selective pressures from viral antagonists, and we show differential responses to vaccinia K3 amongst the duplicates. This highlights the continual innovation occurring in PKR homologs, including species belonging to Haplorhini, the same suborder as humans.

### Tarsiidae Possesses an Ancestral Quadruplication of the Protein Kinase R Gene

Currently, there are six genome assemblies of the Tarsiidae family, spanning four species: *Carlito syrichta* (formerly *Tarsius syrichta*), *Cephalopachus bancanus*, *Tarsius lariang*, and *Tarsius wallacei* (Supplementary Table S1). *Carlito syrichta* represents the reference genome for this primate family and has a predicted annotation using NCBI Gnomon. BLAST homology searching of PKR revealed that there are two truncated and inverted copies upstream of a full-length *PKR* in *C. syrichta*, after which the contig (NW_007255926.1) ends. We refer to the full-length copy as *PKR1* and the additional copies as *PKR2* and *PKR3* (Figure 1A). The truncations in these genes precisely remove the dsRBDs, leaving the kinase domain intact. A third truncated copy, *PKR4*, is found at the start of another contig (NW_007232174.1) (Figure 1A). We considered that this was likely a local quadruplication of the innate immunity factor gene, given that *PKR1-3* genes were adjacent to *GPATCH11* while *PKR4* was adjacent to *SULT6B1*, which are the genes that flank *PKR* across sequenced primate and mammalian genomes (Perelygin, et al. 2006; Vazquez, et al. 2026).

**Figure 1.**
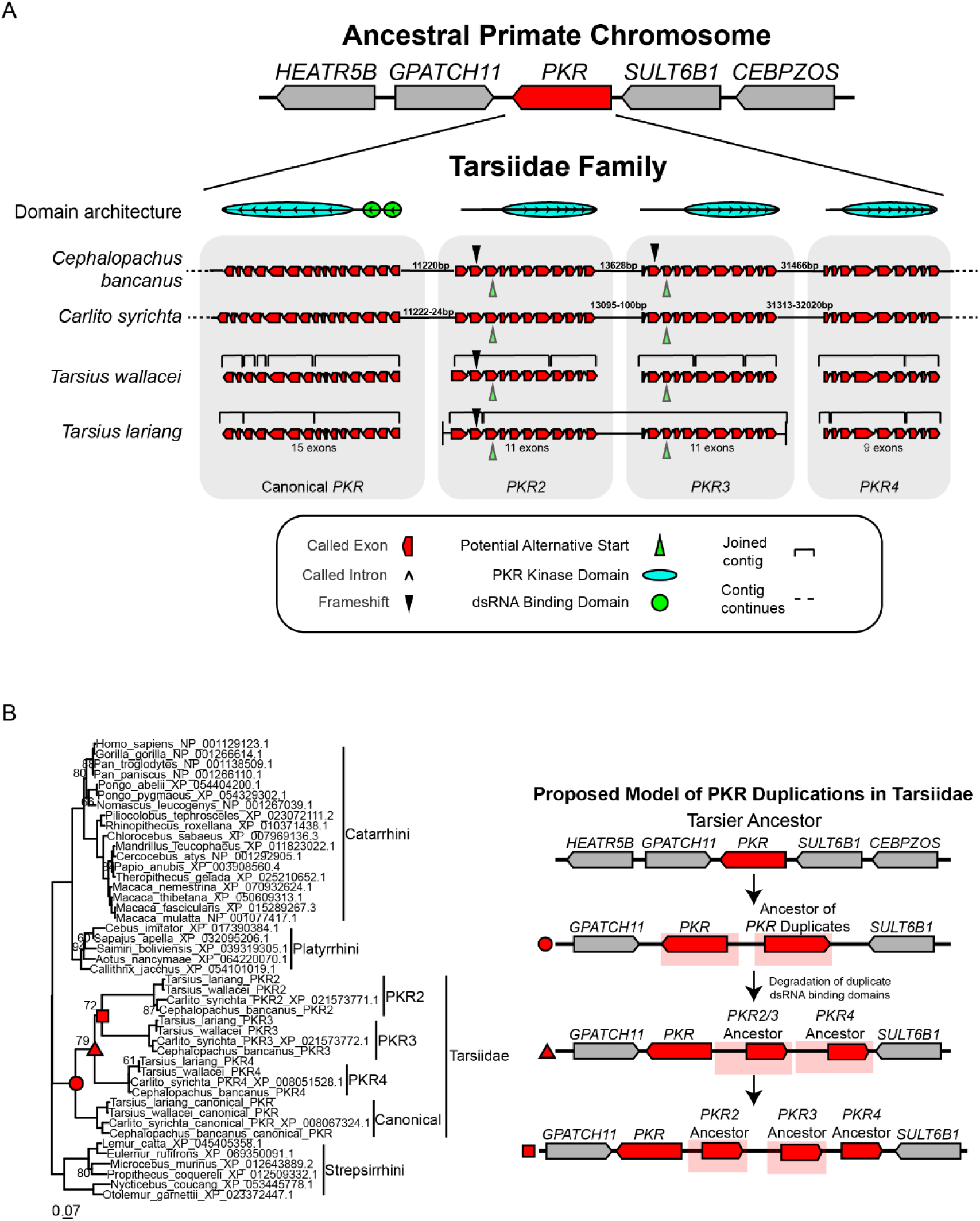
- PKR Quadruplication Across Four Tarsier Species Description of the PKR quadruplication conserved across Tarsiidae. A) Gene structures of the PKR duplicates and their domain architecture across Tarsiidae, with the called exons of each gene depicted to scale. Called exons are angled in the direction of transcription. The ranges given for the intergenic distances for *Carlito syrichta* reflect variation between the reference genome (Schmitz et al. 2016) and a phased-haplotype, long-read assembly (JCASOB000000000.1). The genome assemblies of the *Tarsius* species were particularly fragmented, with several *PKR* called exons scattered across multiple contigs; refer to Supplementary Table 1 for the contigs of these *PKR* genes. B) Maximum likelihood phylogenetic tree of Primate PKR kinase domains rooted at Strepsirrhini. Beside the tree, we outline our proposed model of the order of the PKR duplications in Tarsiidae. All bootstrap values below 95 are shown.

The tarsier *Cephalopachus bancanus* was sequenced recently with PacBio long-read technology (JAPVLO000000000.1), with the resulting assembly resolving this highly repetitive region. All four *PKR* genes are present in this genome and on a single contig, confirming that these three additional copies of *PKR* are indeed an inverted quadruplication of the canonical gene (Figure 1A). During the preparation of this manuscript, a second *Carlito syrichta* long-read genome (PRJNA1484200) was published on NCBI as part of the Vertebrate Genomes Project Phase I (Formenti, et al. 2026), where we also found all four *PKR* genes together in the expected *PKR* syntenic region of primates. Using the predicted exon-intron boundaries from the *Carlito syrichta* annotation, we searched the remaining Tarsiidae genomes and found all four copies in the *Tarsius lariang* and *Tarsius wallacei* genomes, indicating that the quadruplication occurred prior to the split of all present-day tarsier species, and has been maintained since then (Figure 1A; Supplementary Table S1). Across all tarsier genomes, the repeat unit of the quadruplication solely captures the *PKR* gene. The presence of these copies in all four tarsiers dates the Tarsiidae PKR quadruplication between 20 million years (My) old and 58 My old, corresponding to the estimated divergence of crown (extant) tarsiers from each other and the estimated divergence of tarsiers from other primates, respectively (Chatterjee, et al. 2009; Shekelle, et al. 2010; Schmitz, et al. 2016). In agreement with their considerable age, we note that the PKR copies have diverged substantially from one another (Supplementary Figure 1), with no pair sharing more than 70% amino acid identity in their kinase domains. This is equivalent to the amino acid identity shared between that of human and Strepsirrhini PKR kinase domains (95% confidence interval 68.2–81.2 My; Pozzi, et al. 2014).

All of the canonical *PKR1* copies are composed of 15 exons across Tarsiidae, as found across the majority of mammals (Perelygin, et al. 2006). In the *C. syrichta* reference annotation, *PKR2* and *PKR3* have 11 exons while *PKR4* has 9 exons (Figure 1A). We note that in the other three sequenced tarsier species, *PKR2* has an insertion in exon 2 that frameshifts the coding sequence relative to the predicted *C. syrichta PKR2*. Similarly, the sequence of *PKR3* in *C. bancanus* has a frameshift in its exon 2. We believe that the 5′ end of the gene models of the *C. syrichta PKR2* and *PKR3* genes may be incorrect, with conserved downstream methionines being the true start codons (Figure 1A). First, the amino acid composition of the first two exons of *PKR2* and *PKR3* bear little or no sequence homology to any proteins in the BLAST database. Secondly, the lengths of the predicted exons of these duplicates have considerable variation before reaching their kinase domain exons (Supplementary Fig. 2). Taken together, this suggests the ancestor of *PKR2* and *PKR3* would not have encoded dsRBDs and had 9 exons in the Tarsiidae ancestor, as is seen for *PKR4*, if we consider the downstream methionines as the true start codons. According to the phylogeny (Figure 1B), we propose *PKR4* represents the first duplicate to have been maintained in the Tarsiidae ancestor. The ancestor of *PKR2-4* likely lost the exons encoding the dsRNA binding domains, resulting in the subsequent duplicates maintaining a 9-exon gene.

### PKR4 Maintains Kinase Function without the Canonical dsRNA-Binding Domains and Is Resistant to Vaccinia K3

To determine if the additional PKR copies were functional, we used a yeast growth assay in which PKR expression inhibits yeast growth through autoactivation and phosphorylation of the endogenous yeast eIF2ɑ (Dar, et al. 2005). Additionally, we considered both the longer and shorter copies of *PKR2* and *PKR3*, referring to them with an L or S suffix, respectively, to assess if a downstream methionine could be chosen without impacting kinase activity. Importantly, previous work has shown that perturbations of the dsRBDs of human PKR removes its inhibition of yeast growth (Taylor, et al. 2001; Zhang, et al. 2001), indicating that the dsRBDs are required for human PKR’s autoactivation in this assay. Consistent with this result, expression of *C. syrichta* PKR1 (*Cs*PKR1) is toxic, while expression of the dsRBD-lacking *Cs*PKR2 and *Cs*PKR3 displays no kinase activity (Figure 2A) in either the long or short form.

**Figure 2.**
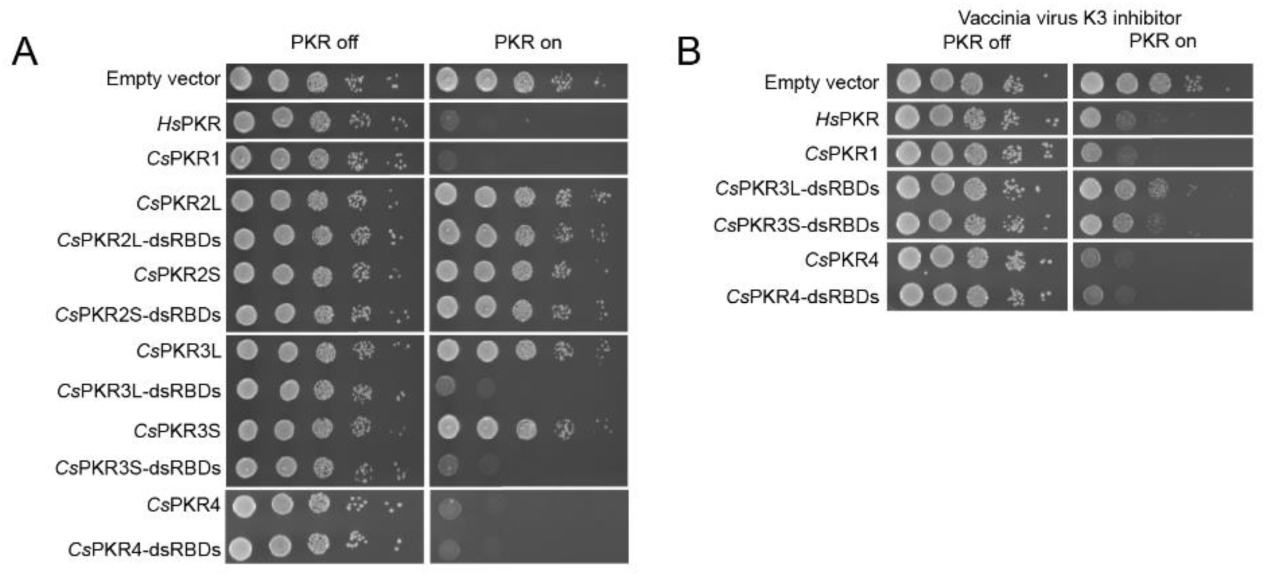
- **Quadruplicated PKRs display diverse functional phenotypes.** Yeast growth assay with *C. syrichta* PKR (*Cs*PKR) copies reveals diverse functionality across the quadruplication. *Homo sapiens* PKR abbreviated *Hs*PKR. *Cs*PKR copies with fusions to the *Cs*PKR1 double-stranded RNA binding domains indicated with “-dsRBDs.” (A) Assay of *Cs*PKR toxicity to yeast. In addition to a plasmid with the tested *Cs*PKR construct, strains contain an empty vector plasmid without the vaccinia virus K3 inhibitor. (B) Extent of inhibition of toxic *Cs*PKRs by the poxvirus pseudosubstrate antagonist vaccinia virus K3.

Interestingly, *Cs*PKR4 displays strong toxicity despite its lack of the dsRBDs (Figure 2A). To assess kinase activity in PKR2 and PKR3, dsRBDs of the canonical CsPKR1 were added to see if this would induce toxicity. Additionally, dsRBDs were added to PKR4 to assess if this would impact toxicity. While the addition of the dsRBDs to *Cs*PKR2S or *Cs*PKR2L does not affect yeast growth, dsRBD addition to both *Cs*PKR3S and *Cs*PKR3L leads to robust toxicity (Figure 2A). Thus, PKR1, PKR3 and PKR4 all retain kinase functionality; PKR2 may have lost kinase function, or it may have retained kinase function that is not autoactivated in yeast by the addition of dsRBDs.

The expansion of a gene in a host may serve to combat pathogenic threats and has previously been hypothesized for the PKR expansions seen in *Myotis* species (Daugherty, et al. 2016; Vazquez, et al. 2026). Many poxviruses carry orthologs of the vaccinia virus pseudosubstrate inhibitor K3 (Elde, et al. 2012; Cao, et al. 2020; Chambers, et al. 2026). We determined if the functional tarsier PKR constructs are differentially inhibited by vaccinia K3 (Figure 2B). The toxicity to yeast growth from *Cs*PKR1 and both functional *Cs*PKR3 constructs (*Cs*PKR3S-dsRBDs and *Cs*PKR3L-dsRBDs) is robustly inhibited, while the toxicity of both *Cs*PKR4 constructs (*Cs*PKR4 and *Cs*PKR4-dsRBDs) is only weakly inhibited (Figure 2B).

### The Tandem *PKR* Genes Conserve Phosphosites and ATP-Binding Sites Necessary for Innate Immune Function

To understand patterns of conservation within the Tarsiidae PKRs, we collected the phosphosites, catalytic site residues, and ATP-binding site residues of human PKR (Uniprot: P19525) that were conserved across primates and identified whether the Tarsiidae PKRs maintain the same amino acid residues (Figure 3) (Consortium 2025). We identified five phosphosites perfectly conserved among full-length primate PKRs (numbered by the human PKR sequence): T258, Y293, T446, T451, and S456. These include T446 and T451, which are in the PKR activation loop and undergo autophosphorylation during PKR activation, whereas T258 and Y293 are noted to either be phosphorylated or be essential for autophosphorylation (Taylor, et al. 1996; Romano, et al. 1998; Taylor, et al. 2001; Su, et al. 2006; Dephoure, et al. 2008). All five sites were preserved in the four sequenced tarsier PKR2-4 orthologs.

**Figure 3.**
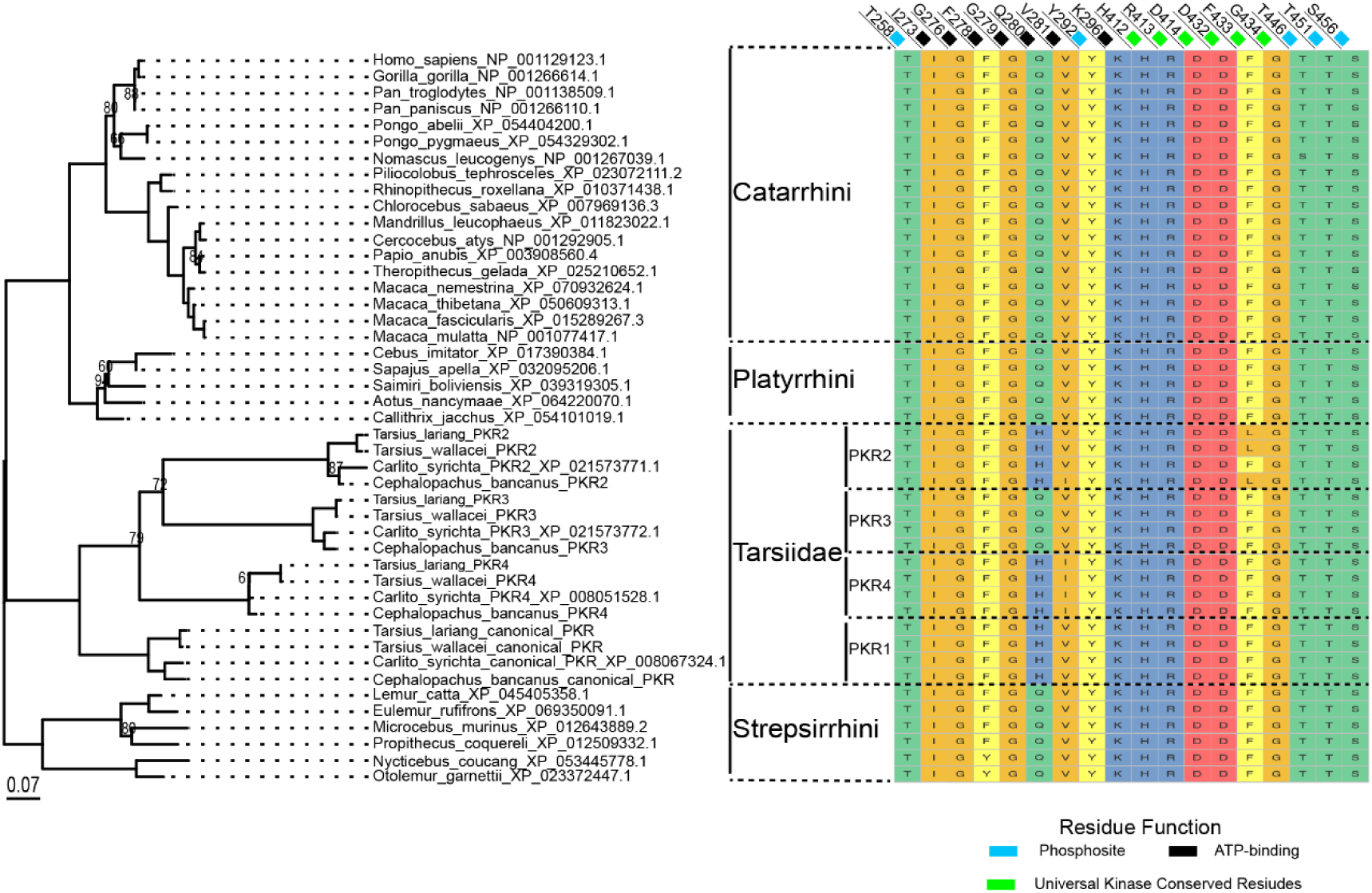
- Kinase Residue Conservation Amongst Tarsier PKR Expansion Lemur-rooted (Strepsirrhini) maximum-likelihood phylogenetic tree of PKR kinase domains with annotation of the residue identities at conserved phosphosites and ATP-binding sites relative to human PKR. The canonical Tarsier PKR has been labelled ‘PKR1’. Bootstrap values below 95 are shown on the phylogenetic tree.

Notably, PKR2 in *C. bancanus*, *T. lariang*, and *T. wallacei* (but not *C. syrichta*) have F433L substitutions in the middle phenylalanine of the catalytic DFG motif, producing a DLG motif like that seen in noncatalytic ROR pseudokinases (Gentile, et al. 2011; Kung and Jura 2019). Otherwise, the catalytic HRD and DFG motifs were perfectly conserved in PKR2-4. Of residues involved in PKR’s ATP-binding, I273, G276, G279, V281, and K296 were preserved in the tarsier PKR2-4 orthologs, aside from a mutation in PKR4 and *C. bancanus* PKR2 of V281 to isoleucine, a chemically similar amino acid substitution that is also present in *Ochotona* (pika) PKRs (XP_040849008.1 and XP_058523342.1). Q280 appears to have mutated to histidine in the pre-duplication ancestral tarsier PKR, which reverted to Q280 in PKR3. We also observed T292I, V298I, and A311T mutations in the *C. syrichta* PKR2 at sites otherwise conserved across primate PKR homologs (Supplementary Fig. 3), but not maintained in other vertebrates (Rothenburg et al. 2009). Overall, in addition to finding expected conservation of functional sites among PKR3 and PKR4, whose kinase domains could induce yeast growth arrest, functional sites were also largely maintained in PKR2, despite its loss of the catalytic DFG motif in three tarsiers and its lack of observed kinase activity in the remaining one. This pattern of conservation is reminiscent of the retention of ATP-binding seen in pseudokinase domains with noncatalytic regulatory function (Zeqiraj, et al. 2009; Hammarén, et al. 2015), which is thought to be important for adopting the conformation of an active kinase. Similarly, if PKR2 needs to achieve an apparent active kinase structure for a noncatalytic role in binding other proteins, that could select for the maintenance of substrate-binding sites and phosphosites.

### Tarsiidae PKRs Have Sites Under Positive Selection Not Found in Other Primates

While functionally constrained residues were conserved in the Tarsiidae PKRs, the PKR expansion and differential inhibition by K3 were consistent with diversification driven by an evolutionary arms race with a virus. We examined if the evolutionary history of tarsier PKRs left a signature of rapid evolution through site-specific enrichment of nonsynonymous mutation. We also investigated if tarsiers contribute more to our understanding of positively selected residues across primates and if there are sites specifically under positive selection within the Tarsiidae family. HyPhy-FEL and the PAML package identified sites of positive selection within Tarsiidae that are similar to sites of selection found across primates (Supplementary File 1 and 2), indicating that the evolutionary pressures faced by tarsier PKRs had commonalities with those faced by other primate PKRs. Interestingly, analyzing non-tarsier primate PKRs versus all primate PKRs, we found that excluding tarsiers from the dataset led to the loss of detection of four additional positively selected residues in both HyPhy-FEL and PAML (HyPhy p<0.05, codeml BEB P>0.95): K426, R447, K493, R534 (Figure 4A; HyPhy-FEL p<0.05, codeml BEB M8 P>0.95). These residues are not conserved within or between the primate suborders, showing that their detection following the inclusion of the Tarsiidae data reveals potential subtle biological signals found across primate evolution (Supplementary Figure 4). K426, R447, K493, and R534 have not previously been identified as positively selected residues in site selection tests of PKR and other ISR kinase domains across primates, but K493 has been identified when comparing PKRs across tetrapods (Elde et al. 2008; Rothenburg et al. 2009). Notably, K426, R447, and K493 are part of the PKR binding interface with eIF2ɑ, and K493 has been implicated as a K3-resistance site (Chambers et al. 2024).

**Figure 4.**
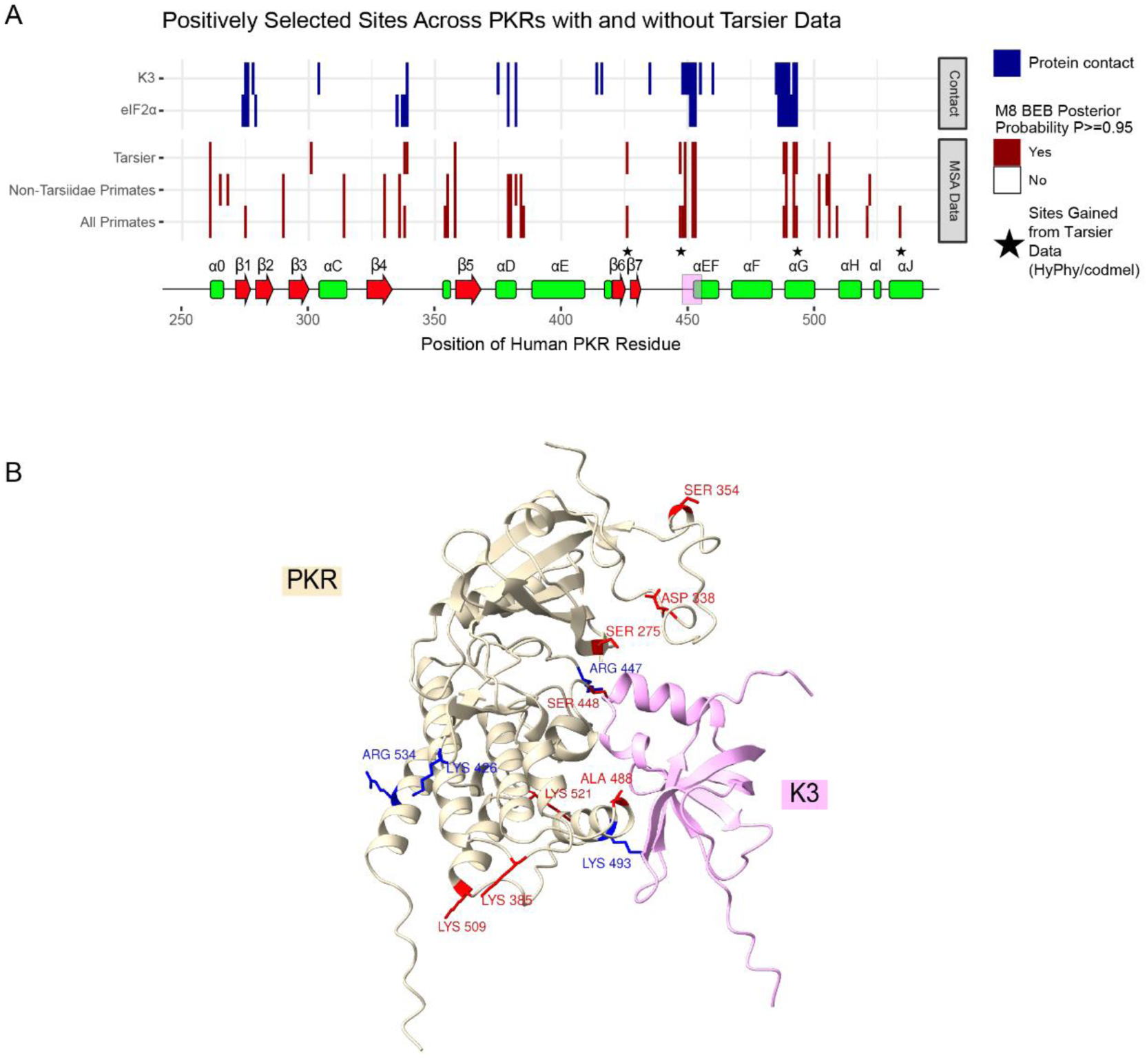
- Sites of Positive Selection in Tarsier PKRs A) We visualized the sites of contact between PKR and eIF2ɑ and PKR and K3 across the kinase domain. We saw several sites of positive selection that aligned at or closely with the protein contact residues. Four residues were shown to gain significance as positively-selected sites between HyPhy-FEL and codeml M8 when the Tarsier data was included in the primate dataset. Below the protein contact and positive selection plot is a schematic of the secondary structure of PKR, to-scale, with the canonical names of the alpha-helices and beta-strands. We have highlighted the activation loop in pink. B) Shown is a superposition of vaccinia virus K3 (1LUZ) onto eIF2ɑ (from 2A1A) to show the K3 interaction interface with PKR (from 2A1A) as described in (Chambers *et al*. 2024), where we have labelled sites in-agreement as positively selected between HyPhy-FEL and codeml M8 (blue) and additionally identified positively selected sites from codeml M8 (red). The majority of positively selected sites are found along the PKR binding interface.

We also found evidence of sites under positive selection within the Tarsiidae lineage using branch-site analysis (branch-site Model A), which survive multiple testing when considering the individual PKR copy lineages: residues N301, S418, and C485 (human numbering) (Bonferroni-corrected p=0.0022, Supplementary Document). S418 and C485 fall within the eIF2ɑ-binding interface, which may be used to evade viral mimics of eIF2ɑ, whereas N301 does not have any clear role in PKR activity or viral mimic interactions. Notably, N301 falls into the positively selected site-class (P>0.95) when only considering tarsier PKR kinase domains. Overall, tarsier PKR data provide statistical power for identifying previously unidentified residues under positive selection and have residues under positive selection that are unique to their lineage.

## Discussion

We have described the only known case of a protein kinase R expansion in primates. Furthermore, this is the only example of an ISR kinase copy number variation that is conserved across an entire infraorder of mammals. We find that the tarsier expansion of PKR involved copies lacking the dsRBDs. Tandem repeats of PKR lacking the dsRBDs have never been confirmed in mammals; the few cases of such PKR homologs appear to be due to genome assembly artifacts from short-read sequencing, later resolved via long-read sequencing (Wu, et al. 2024; Vazquez, et al. 2026). Outside of mammals, PKR genes that do not encode dsRBDs have been identified in cartilaginous fishes (*PKR-sis* genes), and PKR genes encoding Z-DNA binding domains instead of dsRBDs have been found in several fishes (*PKZ* genes) (Rothenburg, et al. 2005; Rothenburg, et al. 2008; Wu, et al. 2020; Wu, et al. 2024). This discovery aligns with previous genomic analyses showing Tarsiidae possess enormous genetic diversity compared to other primates (Schmitz et al. 2016). The diversity provided by this infraorder will continue to greatly improve our understanding of primate biology and evolution.

We have shown that at least 3 of these PKR copies, PKR1, PKR3, and PKR4, have functional kinase domains. Lack of detection of activity from the *C. syrichta* PKR2 upon adding the PKR1 dsRBDs in both the short and long PKR2 forms suggests it may function as a noncatalytic pseudokinase, or it may be a pseudogene. The lack of PKR2 activity cannot be directly attributed to issues in the conserved phosphorylation, ATP-binding, and universal kinase residues given these are identical to the canonical PKR, though a catalytically essential phenylalanine was lost in the other three tarsier PKR2 orthologs. However, mutations at other residues could result in this lack of activity (Supplementary Figure 3). Alternatively, our assay may have lacked the sensitivity to detect any potential activity, or PKR2 was not activated in the context of our assay. In contrast, PKR3 appears to have retained kinase functionality and, as expected, lacks the ability to independently induce activation, rescued by the addition of the dsRBDs.

While it seems unlikely PKR4 would be constitutively active in the native tarsier environment, as this would lead to cell death via the ISR (Zhang, et al. 2001), PKR4 could be regulated at another molecular level, such as through RNA or protein expression levels, or there could be a suppressor of its toxicity present in tarsiers absent from yeast. The constitutive phosphorylation of eIF2ɑ by PKR4, despite lacking dsRBDs, is in contrast to the PKR-sis proteins, where a GST-tag needed to be added for dimerization for their toxicity (Wu, et al. 2024). PKR4, therefore, may have some ability to dimerize despite lacking dsRBDs. The β4-β5 linker has been reported to cause aggregation of human PKR when overexpressed (Dar, et al. 2005). Interestingly, the β4-β5 linker of PKR4 is substantially longer than that of other primates, where its length is 55 or 62 amino acids compared to the ∼26 amino acid average of non-Tarsiidae primate species (Supplementary Figure 3). Alternatively, PKR4’s robust activation may be a feature of the yeast growth assay itself, if there is some activating trigger that is absent from the uninfected native environment.

While these constructs were expressed and tested individually, they have the potential to physically interact with each other in the native environment, which could influence their activation patterns. Indeed, fragments of human PKR can be phosphorylated *in vitro* by full-length PKR, so it is possible that the canonical PKR in tarsiers may interact with these copies (Taylor, et al. 2001). In particular, trans-autophosphorylation of PKR3 by PKR1 following dsRNA binding could allow PKR3’s functional kinase domain to phosphorylate eIF2α, and thus any viral inhibitor that directly binds PKR would need to successfully interface with two highly diverged PKR homologs. We are interested in examining PKR quadruplication co-expression to understand how the copies may interact with one another and provide more insight into their natural biological activity.

The expansion of PKR in tarsiers is a striking mirror to expansions of K3 orthologs in Poxviridae, which occur in an accordion-like fashion (Elde, et al. 2012; Sasani, et al. 2018). Poxviruses use this accordion strategy to quickly adapt to host PKRs (Elde, et al. 2012). The PKR expansion could have been initiated via an initial FoSTeS/MMBIR mechanism, causing the initial inversion duplication (Figure 1B). The FoSTeS/MMBIR mechanism can lead to such large duplicated inversions (>10kb) while maintaining syntenic gene order, whereas other inverted duplication mechanisms would lead to dicentric chromosomes and chromosome terminal deletions following the duplication target (Lee, et al. 2007; Zhang, et al. 2009; Zuffardi, et al. 2009; Hermetz, et al. 2014). Selection by an expansion of a viral mimic or a variety of viral effectors could be a selective pressure leading to copy number variation of PKR in Tarsiidae. We have demonstrated that many of the positively-selected residues are found in the eIF2ɑ/K3 binding interface of PKR, making viral effectors a potential explanation for this innate immunity factor expansion.

In *Myotis* bats, there is haplotype variation in *PKR* copy number, whereas the recent *C. syrichta* long-read haplotype phasing from the Vertebrate Genomes Project and the conservation across the four sequenced tarsier species suggests the tarsier *PKR* expansion has been stably maintained since the divergence of Tarsiidae. In many cases of copy number variation, a major selective pressure driving the return to original copy number is stoichiometry (dosage) within the impacted pathway, where the expansion of one gene can cause downstream strain on the system leading to disease states (Rice and McLysaght 2017). Given PKR is a self-regulatory protein, this may not necessarily drive a return to a single copy. An additional constraint on the reversion of a CNV, following loss of selective pressure, is if the CNV is caused by aneuploidy or segmental duplication, where aneuploidy is reverted far more consistently than segmental duplications (De, et al. 2026). Overall, this means it is unclear if the current retention of the PKR expansion reflects an ongoing or removed selective pressure. Regardless, the divergence in response to K3 (Figure 2B) and striking positive selection in the PKR binding interface (Figure 4), including tarsier-specific sites, alludes to a potential ancient molecular arms race between tarsier PKRs and viral effectors, with tarsiers utilizing genetic innovations unlike any PKR host-virus interaction previously discussed in primates.

The exact role of the PKR duplicates in tarsiers leaves room for some speculation. It is possible that all duplicates function as decoys to sequester viral mimics, which would explain the maintenance of PKR2, the copy lacking kinase activity. The PKR3 and PKR4 copies, which maintain kinase activity but lack dsRBDs, may have the ability to trans-phosphorylate the canonical PKR, while resisting viral mimics. And, finally, recent evidence suggests that mRNAs involved in the immune system may be chimeric, allowing the swapping of domains from one set of immunity genes with another (Venezia, et al. 2026). Given their similar intron-exon structures, it may be possible that PKR3 and PKR4 mRNAs are spliced with dsRBDs from the canonical PKR mRNA. Expression data from Tarsiidae would shed light on all of these questions and broaden perspectives on primate immunity and evolution.

## Methods

### Plasmids and cloning

*Cs*PKR constructs were synthesized (Twist Biosciences, San Francisco, CA) into the human PKR wildtype plasmid MSp509 (previously described in Chambers et al., 2024) with codons optimized for expression in *Saccharomyces cerevisiae* (Supplementary Table 2) using custom Python (v3.11.16) code. Plasmids MSp517 with vaccinia virus K3, MSp515 empty vector K3 inhibitor, and MSp614 empty vector PKR, were previously constructed (described in Chambers et al., 2026). PKR-containing plasmids had a uracil auxotrophic selection marker with the PKR of interest under the galactose inducible promoter pGAL/CYC1. The K3-containing plasmids contained a leucine auxotrophic selection marker with the K3 of interest under a constitutive pTDH3 promoter.

### Yeast strains

*S. cerevisiae* strain BY4742 (*MAT*α *ura3*Δ0 *leu2*Δ0 *his3*Δ1 *lys2*Δ0; Brachmann et al., 1998) was transformed with plasmid containing a poxvirus pseudosubstrate inhibitor or an empty vector using a standard high efficiency lithium acetate transformation protocol (Gietz and Schiestl, 2007). BY4742 containing a poxvirus pseudosubstrate inhibitor plasmid was subsequently transformed with plasmid containing a PKR construct using a modified Frozen-EZ Yeast Transformation II Kit (Zymo, Catalog #T2001) Micro-Scale Protocol. 100 ng of PKR plasmid DNA was transformed and 20 µL of transformation mixture was streaked onto a CSM-Leu-Ura agar plate supplemented with 2% glucose. The remainder of the protocol was followed per the manufacturer’s guidelines.

### Spotting assay

Strains were grown in CSM-Leu-Ura (Sunrise Science Products, Catalog #1038-100) supplemented with 2% glucose at 30°C with shaking at 200 rpm. The following day, the strains were diluted to an initial 1 OD/mL and then serially diluted tenfold to 0.1, 0.01, 0.001, and 0.0001 OD/mL and spotted onto an agar plate with either 2% glucose or galactose. Plates were incubated at 30°C for 2 days then imaged. Spotting was performed in triplicate and a representative image is shown.

## Data analysis

Primate PKRs were collected using the NCBI Orthologs database (Oh, et al. 2025). InterPro was used to identify the elongation factor 2-alpha kinase 2 domains (PTHR11042) of primate PKR homologs (Thomas, et al. 2022; Blum, et al. 2025). Sequences were aligned using MUSCLE v5.1 (Edgar 2022), and alignment were visualized using Jalview (Waterhouse, et al. 2009). All trees were generated using IQ-TREE v2.3.5 with UF-Boot for 1000 replicates and Modelfinder (Kalyaanamoorthy, et al. 2017; Hoang, et al. 2018; Minh, et al. 2020), and all tree visualizations were made using the treeio and ggtree packages in RStudio (Wang, et al. 2020; Xu, et al. 2022). Site selection tests were performed using codeml models 0, 1, 2, 7, and 8 were performed using unrooted trees as input, Pal2Nal codon alignments of the primate kinase domains, and with the codon model FMutSel to assume unequal selection acting on synonymous codons (Suyama, et al. 2006; Yang and Nielsen 2008; Álvarez-Carretero, et al. 2023). HyPhy-FEL was used as a separate site selection model to identify overlaps in the codeml-identified positively selected sites (Kosakovsky Pond and Frost 2005; Kosakovsky Pond, et al. 2020). Branch-site tests were performed in codeml using model A, comparing the null model (fixed omega in a given Tarsier foreground) to the alternative model (estimating omega>1 in a given Tarsier foreground) (Álvarez-Carretero, et al. 2023). 3D structures of PKR and K3 were visualized and labelled using ChimeraX (Meng, et al. 2023).

## Data availability statement

Code and data are available at: https://github.com/SadhuLab/TarsiidaeStudy

## Supporting information

Supplementary File 1- Codeml Analyses

Supplementary File 2- HyPhyFEL Tables

## Acknowledgements

We thank S Rothenburg, T Dever, M Chambers, and all members of the Sadhu lab for helpful discussion. This work utilized the computational resources provided by the NIH HPC Biowulf Cluster (http://hpc.nih.gov). This work was supported by the Intramural Research Program of the National Human Genome Research Institute, NIH (1ZIAHG200401). The contributions of NIH authors are considered Works of the US Government. The findings and conclusions presented in this article are those of the authors and do not necessarily reflect the views of the NIH or the US Department of Health and Human Services.

## Author contributions

T.R.G designed and performed experiments and wrote the main text.

P.G.H and M.J.S wrote the main text, designed experiments, and performed data analyses.

---

**Supplementary Figure S1.**
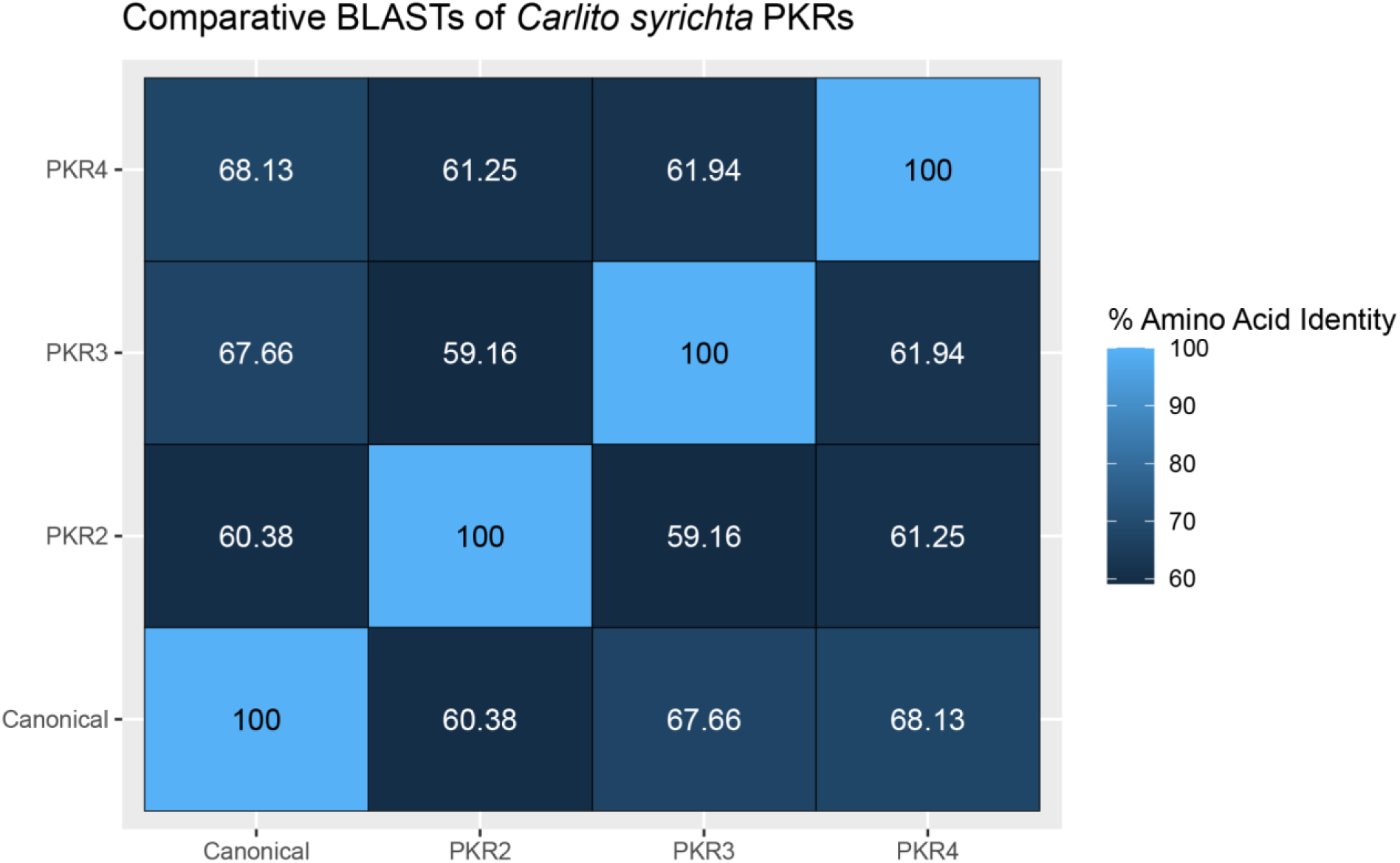
- Pairwise Amino Acid Identity of *Carlito syrichta* PKR paralogs. *Carlito syrichta* PKR paralogs were BLASTed against one another pairwise, with their amino acid identity recorded above.

**Supplementary Figure S2.**
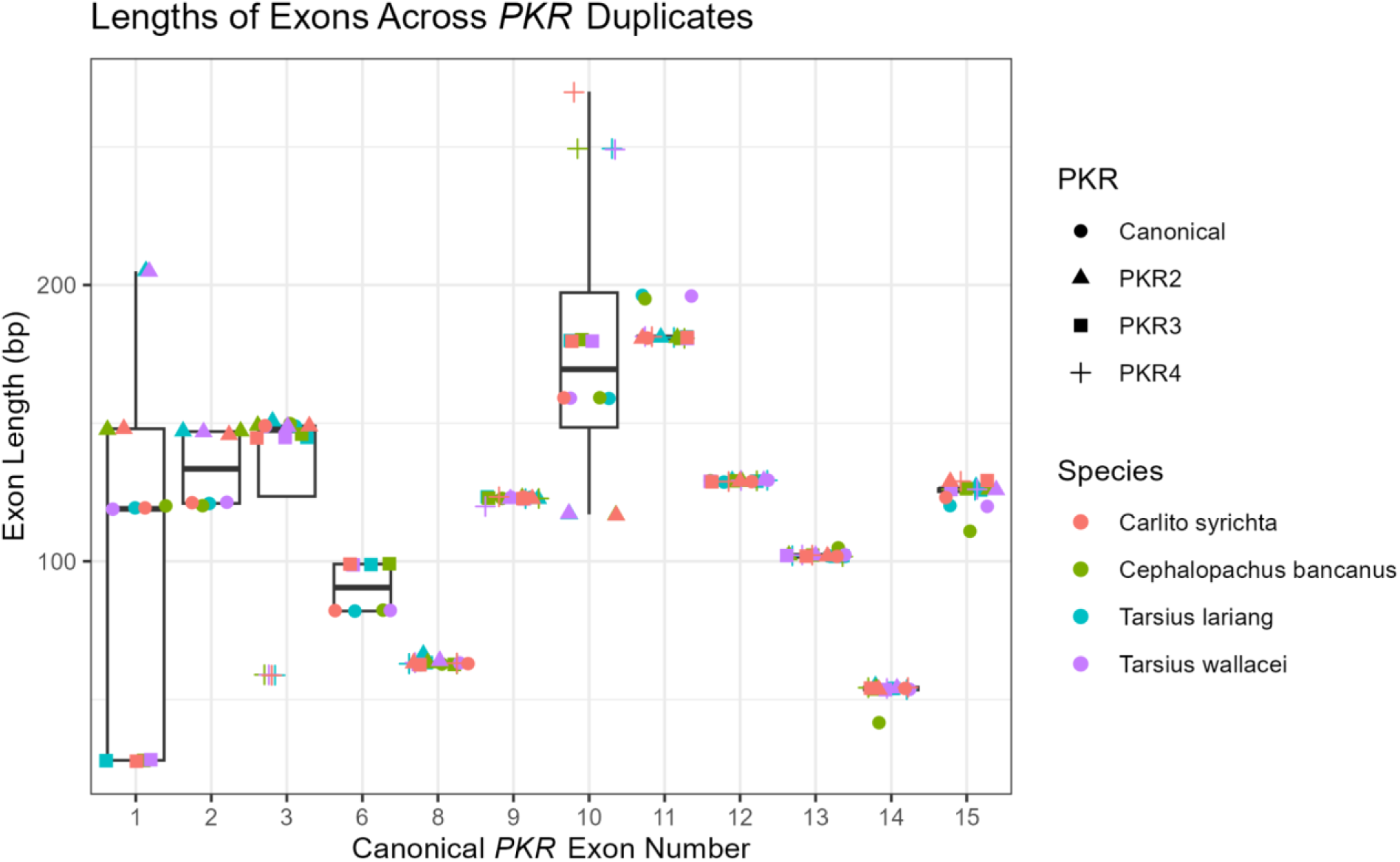
- Exon Length Variation Between PKR Duplicates. The predicted exons of the duplicate *PKR* genes were BLASTed against exons of the canonical *PKR* to understand which canonical exon each duplicate *PKR* exon corresponded to. From here, we plotted the lengths of these predicted exons relative to their canonical exon number. None of the duplicate *PKR* exons were similar to exons 4, 5, or 7 of the canonical gene and are not shown.

**Supplementary Figure S3.**
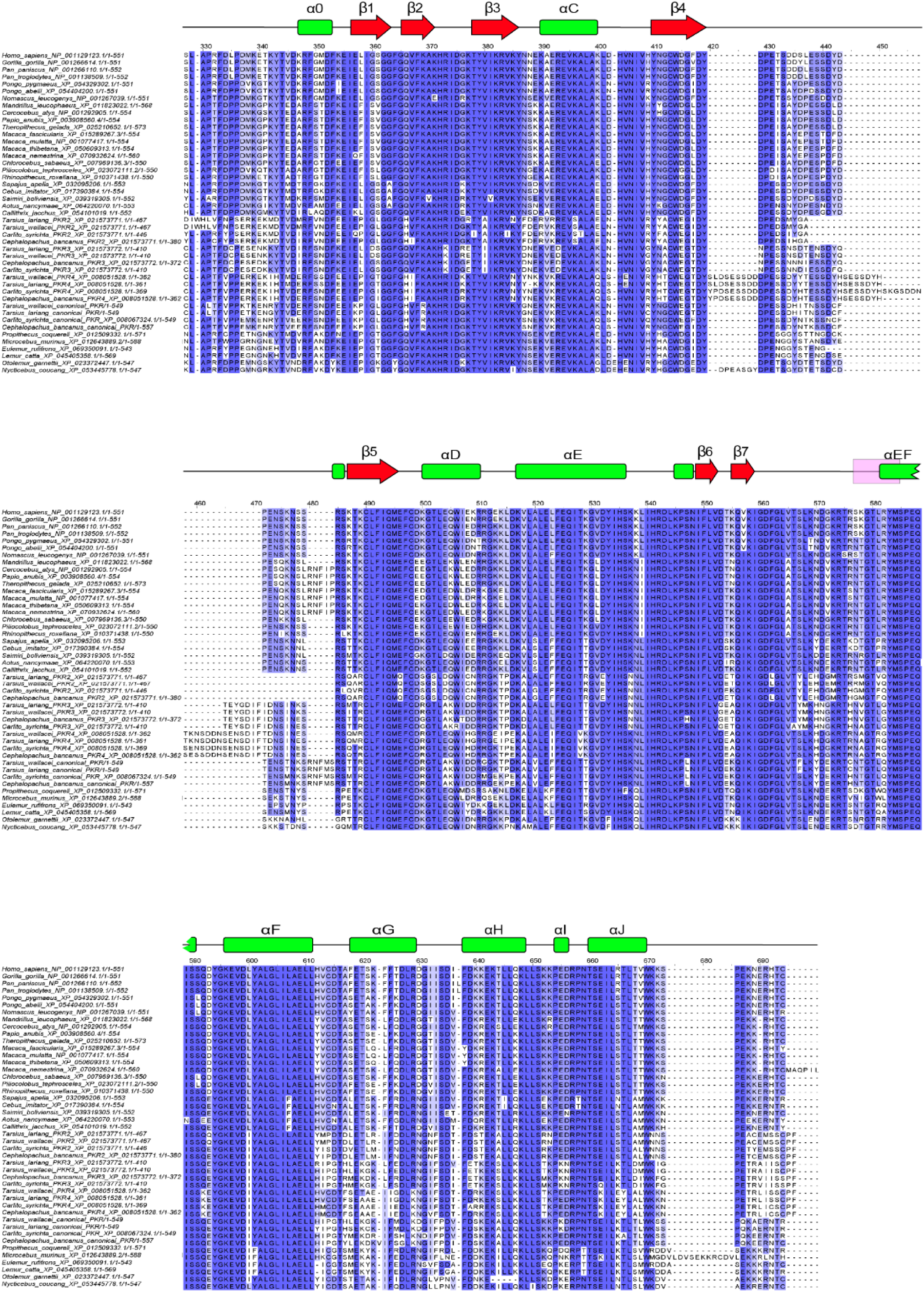
- Full Alignment of PKRs Included In This Study. Protein Kinase R sequences were aligned by MUSCLE and visualized in Jalview (Waterhouse et al. 2009), with intensity of blue corresponding to conservation. The secondary structure of human PKR is shown above the alignment, with the activation loop shaded in pink.

**Supplementary Figure S4.**
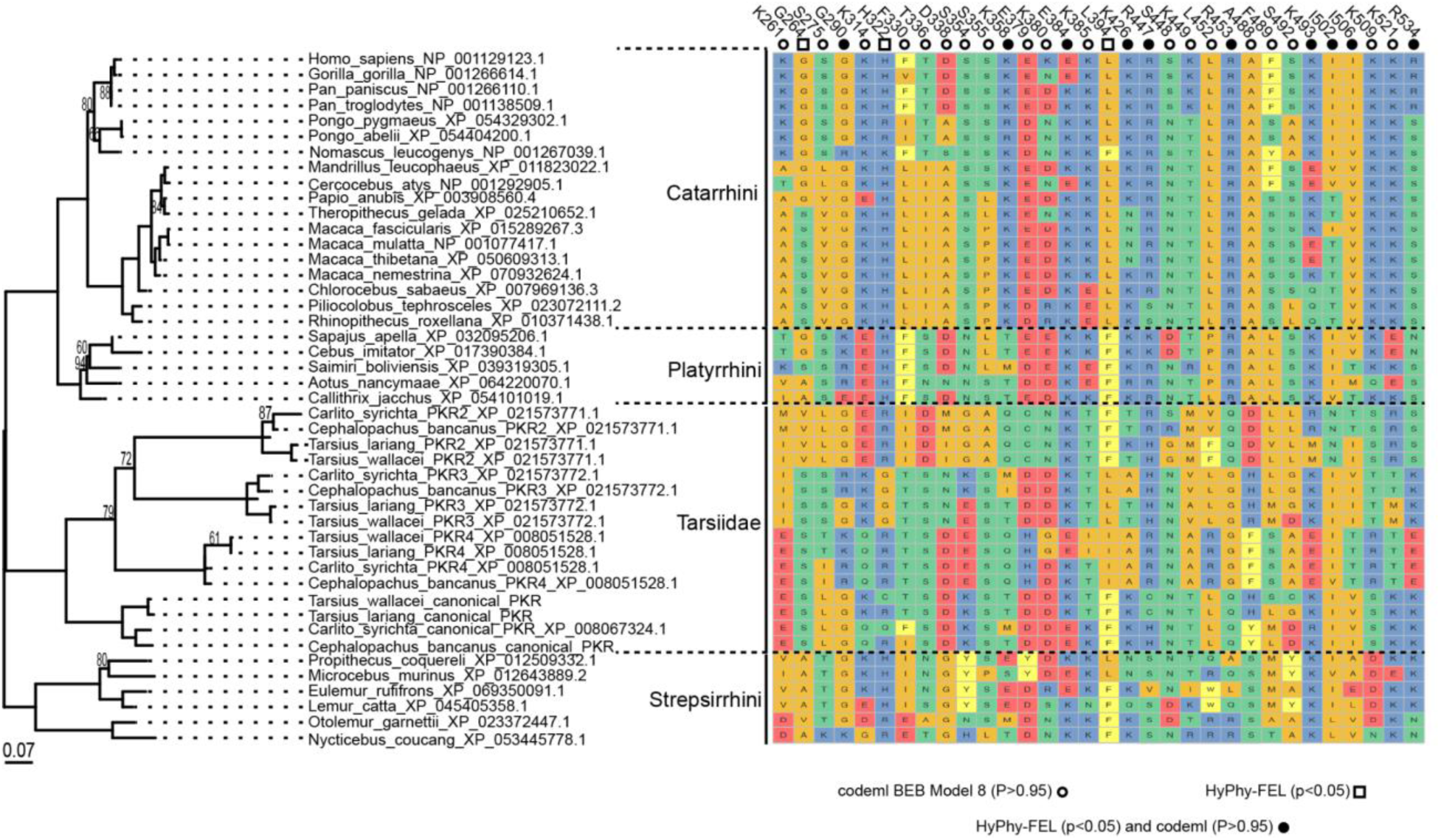
- Positively Selected Sites Across Primate Phylogeny. The phylogenetic maximum-likelihood tree of the PKR kinase domain (left) with the corresponding positively selected residues from each species (right). We have identified where codeml M8 and HyPhy-FEL overlap, as well as sites the models separately identified as positively selected.

