## Supplementary File 1- Codeml Analyses for "Ancestral Quadruplication of Protein Kinase R in Tarsiers Hints Towards Novel Primate-Virus Innovations"

**Supplementary File 1- Codeml Analyses in Heneghan, Grieve, and Sadhu 2026**

**Site Selection Omega Estimates Across All Primate PKR Kinase Domains**

***** Note: the corresponding human residue is shown followed by the posterior Bayes Empirical Bayes probability (P). All P>0.95 shown.

| Model | Likelihood | dN/dS | Estimates of parameters | *Sites with ω > 1 |
| --- | --- | --- | --- | --- |
| M0 | -5862.826189 | 0.74390 | ω= 0.74390 | --- |
| M1a | -5663.037707 | 0.46147 | p0= 0.58409 (p1= 0.41591) ω0=0.07800 | --- |
| M2a | -5606.996897 | 0.84502 | p0= 0.53795, p1= 0.30334 (p2= 0.15872)  ω0=0.08974 | K261 \| 1.000; G290 \| 0.978; K314 \| 0.991; F330 \| 0.998; D338 \| 0.970; S355 \| 0.974; K358 \| 1.000; E379 \| 0.994; K380 \| 1.000; L452 \| 0.999; R453 \| 0.967; F489 \| 1.000; S492 \| 1.000; I502 \| 0.991; I506 \| 1.000 |
| M7 | -5672.260260 | 0.4649 | p = 0.16364, q = 0.18834 | --- |
| M8 | -5607.998454 | 0.8525 | p0 = 0.80144, p = 0.27489, q = 0.49854, (p1 = 0.19856), w = 2.85912 | K261 \| 0.99999; S275 \| 0.99309; G290 \| 0.99717; K314 \| 0.99851; F330 \| 0.9996; T336 \| 0.98464; D338 \| 0.99652; S354 \| 0.97669; S355 \| 0.99578; K358 \| 0.99999; E379 \| 0.99899; K380 \| 0.99999; E384 \| 0.98006; K385 \| 0.96715; K426 \| 0.98996; R447 \| 0.99247; S448 \| 0.96328; K449 \| 0.99027; L452 \| 0.99962; R453 \| 0.99552; A488 \| 0.95875; F489 \| 0.99996; S492 \| 0.99999; K493 \| 0.99052; I502 \| 0.99849; I506 \| 0.99999; K509 \| 0.96706; K521 \| 0.97478; R534 \| 0.97088 |

**All Primate Site Selection Likelihood-Ratio Tests**

| Model | Likelihood Difference | Free parameter | df | 2*deltaL | 𝜒^2^ P-value |
| --- | --- | --- | --- | --- | --- |
| M0 v. M1 | L0= -5862.826189, L1= -5663.037707 | M0=92 v M1=93 | 1 | 399.576964 | 6.808076e-89 |
| M1 v. M2a | L0= -5663.037707  L1= -5606.996897 | M1=93 v M2=95 | 2 | 112.08162 | 4.589712e-25 |
| M7 v. M8 | L0= -5672.260260, L1= -5607.998461 | M7=93 v M8=95 | 2 | 128.523598 | 1.234398e-28 |

**Non-Tarsiidae Primate Site Selection Omega Estimates**

| Model | Likelihood | dN/dS | Estimates of parameters | Sites with ω > 1 |
| --- | --- | --- | --- | --- |
| M0 | -3791.825445 | 0.72885 | ω= 0.72885 | --- |
| M1a | -3676.684425 | 0.42538 | p0= 0.60530 (p1= 0.39470) ω0= 0.05068 | --- |
| M2a | -3641.973319 | 0.84062 | p0= 0.60963, p1= 0.20569 (p2= 0.18468)  ω0=0.08730 | K261 \| 1.000; G290 \| 0.960; K314 \| 0.975; F330 \| 0.988; S355 \| 0.996; K358 \| 0.997; E379 \| 0.986; K380 \| 1.000; L452 \| 0.983; F489 \| 0.990; S492 \| 0.970; I502 \| 0.994; I506 \| 1.000 |
| M7 | -3680.382083 | 0.44050 | p = 0.01413, q = 0.01797 | --- |
| M8 | -3643.247252 | 0.86689 | p0 = 0.76703, p = 0.33184, q = 1.02834, (p1 = 0.23297), w = 2.93000 | K261 \| 0.99995; M265 \| 0.99212; K268 \| 0.95787; G290 \| 0.9944; K314 \| 0.99621; F330 \| 0.99816; T336 \| 0.99104; S355 \| 0.99921; K358 \| 0.99952; E379 \| 0.99766; K380 \| 0.99999; R382 \| 0.97533; E384 \| 0.97852; K449 \| 0.96327; L452 \| 0.99714; R453 \| 0.97084; F489 \| 0.99867; S492 \| 0.99529; I502 \| 0.99897; D505 \| 0.9684; I506 \| 0.99994; K522 \| 0.95324 |

**Non-Tarsiidae Primate Site Selection Likelihood-Ratio Tests**

| Model | Likelihood Difference | Free parameter | df | 2*deltaL | 𝜒^2^ P-value |
| --- | --- | --- | --- | --- | --- |
| M0 v. M1 | L0= -3791.825445, L1= -3676.684425 | M0=60 v M1=61 | 1 | 230.28204 | 5.174097e-52 |
| M1 v. M2a | L0= -3676.684425, L1= -3641.973319 | M1=61 v M2=63 | 2 | 69.422212 | 7.948843e-17 |
| M7 v. M8 | L0= -3680.382083, L1= -3643.247252 | M7=61 v M8=63 | 2 | 74.269662 | 6.814272e-18 |

**Tarsier-Only Site Selection Omega Estimates**

| Model | Likelihood | dN/dS | Estimates of parameters | Sites with ω > 1 |
| --- | --- | --- | --- | --- |
| M0 | -2768.730634 | 0.76829 | ω= 0.76829 | --- |
| M1a | -2723.577108 | 0.54047 | p0=0.45953 (p1= 0.54047) ω0=0.0000 | --- |
| M2a | -2714.009786 | 0.8616 | p0= 0.45562, p1= 0.34719, (p2= 0.19720)  ω0=0.00000 | A488 \| 0.954; F489 \| 0.970; S492 \| 0.985 |
| M7 | -2724.069301 | 0.5000 | p = 0.00506, q = 0.00500 | --- |
| M8 | -2714.040117 | 0.95734 | p0= 0.77372 (p1= 0.22628)  p=0.00500, q=0.00755, ω=2.47012 | K261 \| 0.98688; N301 \| 0.98293; D338 \| 0.98307; D339 \| 0.9849; K358 \| 0.98796; K426 \| 0.966; R447 \| 0.95209; K449 \| 0.96688; L452 \| 0.98458; R453 \| 0.97214; A488 \| 0.99116; F489 \| 0.99427; S492 \| 0.99746; K493 \| 0.97125; I506 \| 0.98689 |

**Tarsier-Only Site Selection Likelihood-Ratio Tests**

| Model | Likelihood Difference | Free parameter | df | 2*deltaL | 𝜒^2^ P-value |
| --- | --- | --- | --- | --- | --- |
| M0 v. M1 | L0= -2768.730634, L1= -2723.577108 | M0=34 v M1=35 | 1 | 90.307052 | 2.039245e-21 |
| M1 v. M2a | L0= -2723.577108, L1= -2714.009786 | M1=35 v M2=37 | 2 | 19.134644 | 6.997854e-05 |
| M7 v. M8 | L0= -2724.069301, L1= -2714.040117 | M7=35 v M8=37 | 2 | 20.058368 | 4.409413e-05 |

**Estimations of Background and Foreground Omega Values in Branch-Site Models (Model A)**

| **Foreground branch** | **Site class** | **Proportion** | **Background omega** | **Foreground omega** |
| --- | --- | --- | --- | --- |
| **Whole Tarsier Clade** | 0 | P0 = 0.52912 | W0=0.06350 | W2=0.06350 |
|  | 1 | P1=0.30949 | W1 =1.0000 | W2=1.0000 |
|  | **2a** | **P2a=0.10183** | **W0=0.06350** | **W2=2.23981** |
|  | **2b** | **P2b=0.05956** | **W1=1.0000** | **W2=2.23981** |
| Only Canonical PKR | 0 | P0=0.58409 | W0=0.07800 | W0=0.07800 |
|  | 1 | P1= 0.41591 | W1=1.000 | W1=1.000 |
|  | 2a | P2a=0 | W0=0.07800 | W2=1.000 |
|  | 2b | P2b=0 | W1=1.000 | W2=1.000 |
| PKR2 Clade | 0 | P0=0.49546 | W0=0.07428 | W0=0.07428 |
|  | 1 | P1=0.34548 | W1=1.00000 | W1=1.00000 |
|  | 2a | P2a=0.09372 | W0=0.07428 | W2=2.66726 |
|  | 2b | P2b=0.06535 | W1=2.66726 | W2=2.66726 |
| PKR3 Clade | 0 | P0=0.58409 | W0=0.07800 | W0=0.07428 |
|  | 1 | P1=0.41591 | W1=1.00000 | W1=1.00000 |
|  | 2a | P2a=0.0000 | W0=0.07800 | W2=1.00000 |
|  | 2b | P2b=0.0000 | W1=1.00000 | W2=1.00000 |
| PKR4 Clade | 0 | P0=0.58102 | W0=0.07744 | W0=0.07744 |
|  | 1 | P1=0.40949 | W1=1.00000 | W1=1.00000 |
|  | 2a | P2a=0.00557 | W0=0.07744 | W2=2.42803 |
|  | 2b | P2b=0.00392 | W1=1.00000 | W2=2.42803 |

**LRTs of Branch-Site Models (Model A v Model A_null_) Positive Selection Across Tarsier Lineage**

| **Foreground Branch** | **Likelihood** | **Free parameters** | **df** | **2*deltaL** | **P value** | **Bonferroni Correction** | Sites with ω > 1 |
| --- | --- | --- | --- | --- | --- | --- | --- |
| **Whole Tarsier Clade** | **Null= -5651.632760, Alt = -5645.457787** | **Null=94, Alt=95** | **1** | **12.349946** | **4.40e-4** | **0.0022** | N301 \| 0.996; S418 \| 0.963; C485 \| 0.969 |
| Only Canonical Clade | Null = - 5663.037500  Alt = - 5663.037501 | Null=94, Alt=95 | 1 | 0 | 1 | 5 | --- |
| PKR2 Clade | Null = -5661.292254, Alt = -5660.552950 | Null=94, Alt=95 | 1 | 1.478608 | 0.2239924 | 1.119962 | --- |
| PKR3 Clade | Null= -5663.037501, Alt= -5663.037501 | Null=94, Alt=95 | 1 | 0 | 1 | 5 | --- |
| PKR4 Clade | Null=-5663.011285, Alt= -5662.890118 | Null=94, Alt=95 | 1 | 0.242334 | 0.6225254 | 3.112627 | --- |
